# Food texture preference reveals multisensory contributions of gustatory organs in behaviour and physiology

**DOI:** 10.1101/2024.07.04.602043

**Authors:** Nikita Komarov, Cornelia Fritsch, G. Larisa Maier, Johannes Bues, Marjan Biočanin, Clarisse Brunet Avalos, Andrea Dodero, Jae Young Kwon, Bart Deplancke, Simon G. Sprecher

## Abstract

Food presents a multisensory experience, with visual, taste, and olfactory cues being important in allowing an animal to determine the safety and nutritional value of a given substance^1^. Texture, however, remains a surprisingly unexplored aspect, despite providing key information about the state of the food through properties such as hardness, liquidity, and granularity. Food perception is achieved by specialised sensory neurons, which themselves are defined by the receptor genes they express. While it was assumed that sensory neurons respond to one or few closely-related stimuli, more recent findings challenge this notion and support evidence that certain sensory neurons are more broadly tuned. In the *Drosophila* taste system, gustatory neurons respond to cues of opposing hedonic valence or to olfactory cues. Here, we identified that larvae ingest and navigate towards specific food substrate hardnesses, and probed the role of gustatory organs in this behaviour. By developing a genetic tool targeting specifically gustatory organs, we show that these organs are major contributors for evaluation of food texture and ingestion decision-making. We find that ablation of gustatory organs not only results in loss of chemosensation, but also navigation and ingestion preference to varied substrate textures. Furthermore, we show that certain neurons in the primary taste organ exhibit varied and concurrent physiological responses to mechanical and multimodal stimulation. We show that individual neurons house independent mechanisms for multiple sensory modalities, challenging assumptions about capabilities of sensory neurons. We propose that further investigations, across the animal kingdom, may reveal higher sensory complexity than currently anticipated.

## Introduction

The properties of food play a crucial role in an animal’s decision to ingest. While smell, taste, and visual properties provide important details about a food source, texture is a property of food that is additionally critical. The texture of food serves as a multi-dimensional attribute of parameters not obviously determined by other sensory organs. Thus, the mechanical sensation of food sources is necessary for an animal’s ability to completely evaluate the food it encounters.

The full extent of sensory roles of the *Drosophila* larval external sensory organs is not known. For example, although the presence of mechanosensory neurons in the primary taste sensing centre, the terminal organ (TO) was already suggested, identification of mechanisms, responses, and functions has proved elusive ^2,3^. While it has been assumed that mechanosensation is important for decision-making, few studies have been conducted to elucidate the role of peripheral mechanosensation in the larva ^2,4,5^. Meanwhile, the role of mechanosensation as a critical component of food decision-making in adults was recently characterised ^1^.

The perception of external cues is achieved by highly specialised sensory neurons. Different types of sensory neurons are thought to be tuned in a narrow fashion, thereby responding to a defined type of stimulation such as a specific range of wavelength of light or class of chemical compounds. Narrow tuning is assumed to be a critical feature of stimuli coding, allowing tightly regulated processing and integration in defined brain circuits. An essential function of taste systems revolves around distinguishing appetitive and aversive cues (*e.g.,* ‘bitter’ vs. ‘sweet’) at the level of the sensory neuron. Since this is the first point of contact with the chemical cue, a certain amount of debate is present about whether individual neurons can detect unique or multiple modalities. On the one hand, it is believed that neurons are either specifically or broadly tuned to one of 5 canonical tastes – sweet, bitter, umami, sour, and salt ^6–8^. This is referred to as the “labelled-line” model. On the other hand, recent findings uncovered that individual taste neurons of both *Drosophila* larvae and mice are responsive to multiple modalities, including opposite hedonic valence ^9–11^. This indicates that the organisation, coding, and function of the peripheral chemosensory organs are more intricate than previously thought. Furthermore, the concept of an individual neuron, rather than the organ as a whole, integrating other senses such as light, mechanosensation, thermosensation, or hygrosensation has been suggested but remains to be explored ^12^.

The larva of the fruit fly *Drosophila melanogaster* provides a powerful model to uncover mechanisms of sensory perception due to its relative neuronal numerical simplicity, ample genetic tools, and, importantly, traceable processing and stereotypic behavioural responses ^13^. Moreover, the larva represents a highly relevant model for exploring sensory systems and food consumption due to its biological need to ingest as much food of the highest quality possible. Failing to do so, the larva will either not undergo metamorphosis or develop into a smaller adult ^14^. Larval taste is separated into external and internal components. On the exterior, the head of the larva bilaterally houses terminal organs (TO) – the primary taste centre – and the dorsal organs (DO) – the primary olfactory centre. Additionally, the ventral organ (VO) is also believed to be involved in taste, as well as other sensory modalities ^15^. After ingesting food, larvae can taste food using pharyngeal sensilla, located along the oesophagus inside the mouth opening and projecting their dendrites into the gastrointestinal tract. Moreover, larvae are able to sense sugar not only in the sensory organs but also in the brain, where a receptor attributed to fructose sensing is expressed, and this function is attributed to sensing the internal nutritional state of the animal ^16^.

The molecular basis of taste sensing is not fully understood. In the olfactory system, an individual *Odorant receptor* (*Or*) or *Ionotropic receptor* (*Ir*) gene is expressed alongside the obligate *Odorant receptor co-receptor* (*Orco*) or one of two *Ir*-co receptors, respectively. In taste neurons, the organisation is different: *gustatory receptors* (*Gr*), *Irs*, and other putative chemosensors, such as the *pickpocket* (*ppk*) family, are co-expressed in an unclear manner ^15^. Furthermore, the nature of Grs as channel-forming or signal-conducting proteins is not known, in contrast to the resolved stetramerisation of the OR complex in olfactory neurons. One exception is the CO_2_-sensing complex comprised of Gr21a and Gr63a, which together confer carbonation sensing, but not either receptor alone ^17^. Beyond this, a range of receptor genes have been proposed for specific modalities, such as *Gr66a* for bitter sensing or *ppk11* and *ppk19* for salinity ^18–21^. Interestingly, despite sugar being a critical nutritional cue, no peripheral receptor has been identified. The canonical sugar sensor, *Gr43a*, is expressed in the pharyngeal sensilla and in the brain but not in the TO or VO ^16^. Conversely, larvae are able to sense sugar at the periphery through multiple neurons extending their dendrites into the TO, albeit only one of these, C2, has a behavioural phenotype when silenced ^10,11^. While being essential for larval survival and growth, the mechanisms for these responses have not yet been elucidated.

In order to study the role of the gustatory organs, we created a novel split-Gal4 line which drives reporter expression in the peripheral gustatory organs (GO). By using behaviour assays, this tool allows us to demonstrate that these organs contribute not only to taste but also to mechanosensation. Additionally, by employing whole-organ and single-neuron volumetric live imaging, we show that individual neurons respond to chemical and mechanical stimuli. Furthermore, we show that one of these gustatory sensory neurons (GSN) is multimodal, given its responses to both sugar and CO_2_, as well as multisensory and ability to respond to mechanical stimulation. Thus, we propose that multisensory integration in individual neurons may modulate their output, demonstrating a mechanism for context-based responses at the single-neuron level. Hereby, we show that a comparatively simple taste system integrates a significantly larger number of inputs than previously thought, which may account for a maggot’s fascinating ability to distinguish a wide variety of taste stimuli.

## Results

### Larvae navigate to a specific range of substrate hardnesses corresponding to specific stages of fruit decomposition

While it has been reported that larvae prefer softer food substrate textures ^5^, we aimed to determine the range of preference exhibited by freely-behaving animals. This would assess, for instance, whether larvae will navigate to a harder-textured unripe fruit (*e.g.*, apple) compared to a more ripe one (Figure 1A left). Additionally, the texture of food could determine whether an animal will ingest it (Figure 1A right). In order to evaluate traditional agarose-based experimental paradigms, we set out to understand how agarose concentration relates to physical properties of a flies’ assumed natural food source – decaying fruit (Figure 1A’). Here, we tested agarose discs of a variety of concentrations (Figure 1A’’, left), as well as dissected apple, pear, banana, and pineapple fruits into similarly sized disks, and allowed to decompose over 5 days (Figure 1A’, right). We observed that freshly-cut fruits, except banana, are significantly harder than even the highest (2.5%) agarose concentration in terms of the compression modulus (Figure 1A’’, right). Freshly cut banana fruit, however, strongly resembles the softness of 1-1.25% agarose concentration.

**Figure 1:**
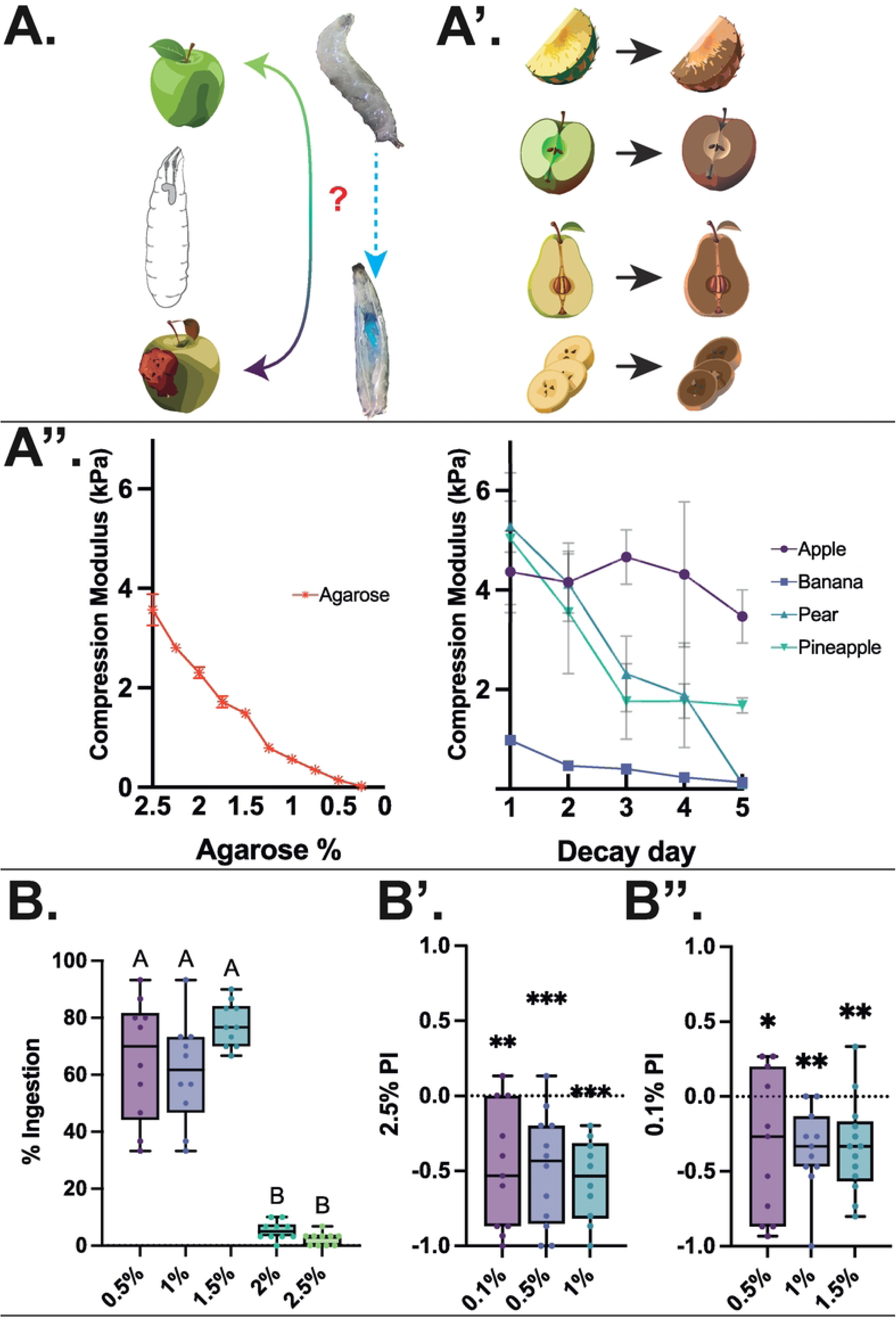
Texture preference in *Drosophila* larvae, as a relevant cue for varying feeding substrates. A: Left – cartoon of larval texture-based environment, showing a harder (fresh) fruit and a softer (ripe) fruit. Right – larval ingestion can be visualised by blue-dyed agarose present and visible in the digestive system. A’: experimental paradigm involving a range of decaying fruit for mechanical analysis. A’’: Mechanical analyses of substrate properties of agarose (left) and decomposing fruit (right) identifying an increasing hardness of agarose directly related to concentration and a decrease in hardness of dissected fruit over time, except for apple until day 5 after sectioning. B: ingestion of a range of agarose concentrations after 2 minutes of exposure. Larvae readily and immediately ingest agarose food substrates up to and including 1.5%, however cease to ingest beyond this threshold. One way Kruskal-Wallis with Dunn’s multiple comparison tests. Different letters denote p<0.05. B’: two-choice navigation preference between 2.5% agarose and 0.1%, 0.5%, and 1% agarose, respectively. Larvae show a consistent preference towards all softer agarose substrates compared to 2.5% agarose. One-Sample t and Wilcoxon test vs 0. **-p<0.01, ***-p<0.001. B’’: two-choice navigation preference between 0.1% agarose and 0.5%, 1%, and 1.5% agarose, respectively. Larvae show a consistent preference for the harder agarose concentrations. One-Sample t and Wilcoxon test vs 0. *-p<0.05, **-p<0.01. N=10-15 trials (x30 individuals) for all behaviour experiments.

Next, we assessed whether larvae will ingest foods of across a variety of hardnesses. Here, we found that larvae readily ingest sucrose-doped blue-dyed agarose substrate at concentrations of 0.5%, 1%, and 1.5% within the first 2 minutes (Figure 1B). However, at 2% agarose concentration and above, larvae almost entirely cease to ingest the substrate. This indicates that there is a specific hardness threshold at which larvae are either unable or unwilling to ingest. Next, we determined whether larvae prefer softer or harder substrates, thus if hardness or softness presents as a specifically aversive or appetitive sensory cue for navigation by means of two-choice assays. Here, larvae were allowed to freely navigate on plates containing two halves of distinct agarose substrates. First, the larvae were given the choice between one half containing 2.5% agarose, and the other half containing one of 0.1%, 0.5%, or 1% agarose. We observed that larvae consistently prefer the softer concentration compared with 2.5% agarose (Figure 1B’). Next, larvae were given a choice between an excessively-soft (0.1%) agarose substrate against 0.5%, 1%, and 1.5% concentrations. Here, we observe that larvae prefer to navigate onto slightly harder agarose (Figure 1B’’). Thus, we observe that larvae appear to prefer the specific softness range between 0.5% and 1.5% agarose, with softer or harder food substrates presenting as aversive stimuli. Therefore, the hardness of the food substrate provides a specific sensory cue that allows the animals to navigate to optimal food textures and preferentially ingest food of this hardness range. This correlates to the hardness of pear and pineapple after 3-4 days of decomposition, highlighting that fruit hardness may provide sensory cues about the state of decomposition.

### Understanding the role of gustatory organs in mechanosensation through the creation of a novel split-GAL4 driver

A basic way to investigate the function of a particular system is to inhibit or ablate it, subsequently observing the resulting phenotype. Thus, in order to investigate the gustatory organs (GOs) specifically, we required a driver that would allow for such manipulations. This would be similar to the role of *Orco*-GAL4 and Orco mutants in the dorsal organ ganglion (DOG), which were used to create anosmic animals effectively ^22,23^. In order to decide on an approach for investing in the sensory system, we needed to understand the molecular profiles of the sensory neurons of the terminal organ ganglion (TOG) and the ventral organ ganglion (VOG) representing the GOs and the primarily olfactory DOG. Organs were dissected, digested, and subsequently sequenced using the <u>d</u>etermin<u>is</u>tic, mRNA-capture bead and cell <u>co</u>-encapsulation dropleting system (DisCo, Bues et al., 2022) (Figure S1). Through analysis of these data using the Seurat package ^25,26^, we isolated filtered objects expressing the neuronal markers *Neuroglian* (*Nrg*), *Synaptobrevin* (*Syb*), *Neuronal Synaptobrevin* (*nSyb*), and *pebbled* (*peb*), resulting in a set of 153 neurons. Moreover, we identified that *Orco*, present in all olfactory cells of the DOG, does not overlap with cells expressing *proboscipedia* (*pb*), a member of the Hox transcription factor family known to mediate the specification of adult mouthparts (Figure 2A, left) ^27^. Using immunofluorescence staining, it emerged that *Pb* inclusively, but not exclusively, labels the neuronal population of the TOG (Figure S1), while it is absent from the DOG. By more specifically targeting sensory neurons inserting the split-GAL4 components into the endogenous loci of the transcription factor Pebbled and of *pb*, we developed a specific split-GAL4 driver for the gustatory organs (Figure 2A right, for details, see materials and methods). Through immunohistochemical stainings and whole-mount live imaging, we determined that the driver covers the GOs but not the DOG (Figure 2B). Thus, we are able to drive reporters of our choosing in the peripheral taste organs specifically, giving us access to the well-established GAL4/UAS toolkit.

**Figure 2:**
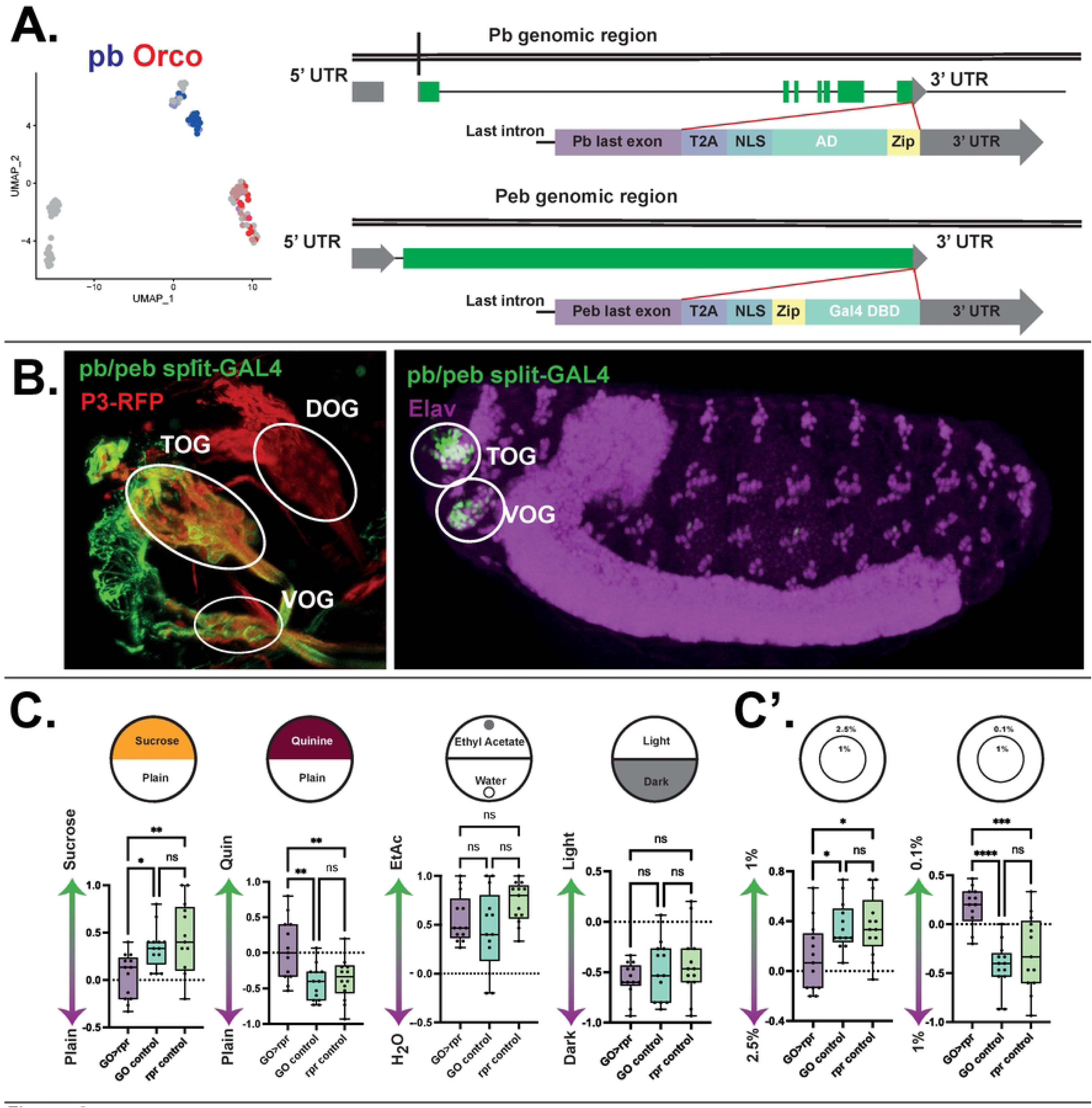
larval taste organ identification, and characterisation of larval taste organ function. A: Left: neuronally-filtered single-cell sequencing expression of the olfactory marker, *Orco*, and *proboscipedia* (*pb*). Right, cloning strategy for creation of a split-GAL4 knock-in. Upper: the activating domain (AD), along with a T2A, NLS and zipper (Zip) domains was inserted between the last exon of Proboscipedia and the 3’ UTR. Lower: the GAL4 DNA binding domain (DBD) was inserted along with T2A, NLS, and Zip domains between the last exon of *pebbled* and the 3’ UTR. B: Immunofluorescence stainings showing the expression of the transgenic P3-RFP and generated *pb/peb*-split-GAL4 driving a UAS-*myrGFP* reporter in the taste organs, but not the DOG (left), With expression in the embryonic phase (right) mirroring that of the larva. C: UAS-*reaper* (*rpr*)-mediated ablation results in defects of appetitive (sucrose) and aversive (quinine) choice. Appetitive olfactory response to Ethyl acetate was unaffected in the ablated condition, as was light-aversive behaviour. C’: UAS-*rpr*-mediated ablation of GOs results in defective substrate hardness preference. Larvae lose preference for the softer 1% agarose versus harder 2.5% agarose, in addition they start to prefer excessively soft 0.1% agarose. N=10-15 for all behaviour assays. One-way ANOVA (Tukey’s multiple comparisons) test. *-p<0.05, **-p<0.01, ***-p<0.001, ****-p<0.0001, ns-Not Significant (p>0.05). Not significant where not shown. Fruit icons were created by Adobe generative fill.

### Identifying the role of the GO contribution to sensory modalities

To uncover whether the broad range of cell types in the GOs results in a role in sensing different environmental stimuli, we selectively ablated neurons expressing both components of the *pb/peb* split-GAL4 by crossing these flies with the pro-apoptotic reporter UAS-*reaper* (*rpr*). Expectedly, we found that in a two-choice behaviour assay, the experimental larvae showed a significant reduction of response to both appetitive (sucrose) and aversive (quinine) agarose (Figure 2C). Additionally, we evaluated whether olfactory preference to an attractive odour (ethyl acetate) or visual aversion to light would be affected and found no significant change (Figure 2C). Thus, we conclude that the GOs do not appear to contribute to olfactory or light sensing. To understand whether the range of proposed mechanosensory neurons of the GOs contributes to mechanical sensing, we used the paradigm of substrate texture preference. Here, we observe that by ablating the GOs, the preference for “soft” substrate (1% agarose) is also significantly reduced, as well as the avoidance of a “very soft” substrate (0.1% agarose) (Figure C’).

### Identification of mechanosensory gene expression in the larval head

Since mechanosensation has been proposed as a feature within the GOs previously, based on neuronal morphology ^3^, we set out to ascertain whether the proposed mechanosensory neurons in the primary sensory organs express canonical mechanoreceptor markers. We probed the scRNAseq dataset and found that three genes involved in mechanosensory functions are present: *nanchung* (*nan*), *no mechanoreceptor potential C* (*NompC*), and *painless* (*pain*). The confidence for the expression was increased by means of immunofluorescence staining, finding 2-3 *nan-* and *NompC*-expressing cells in the larval head, along with a relatively broad expression of *pain* (Figure S2, Figure 3A).

**Figure 3:**
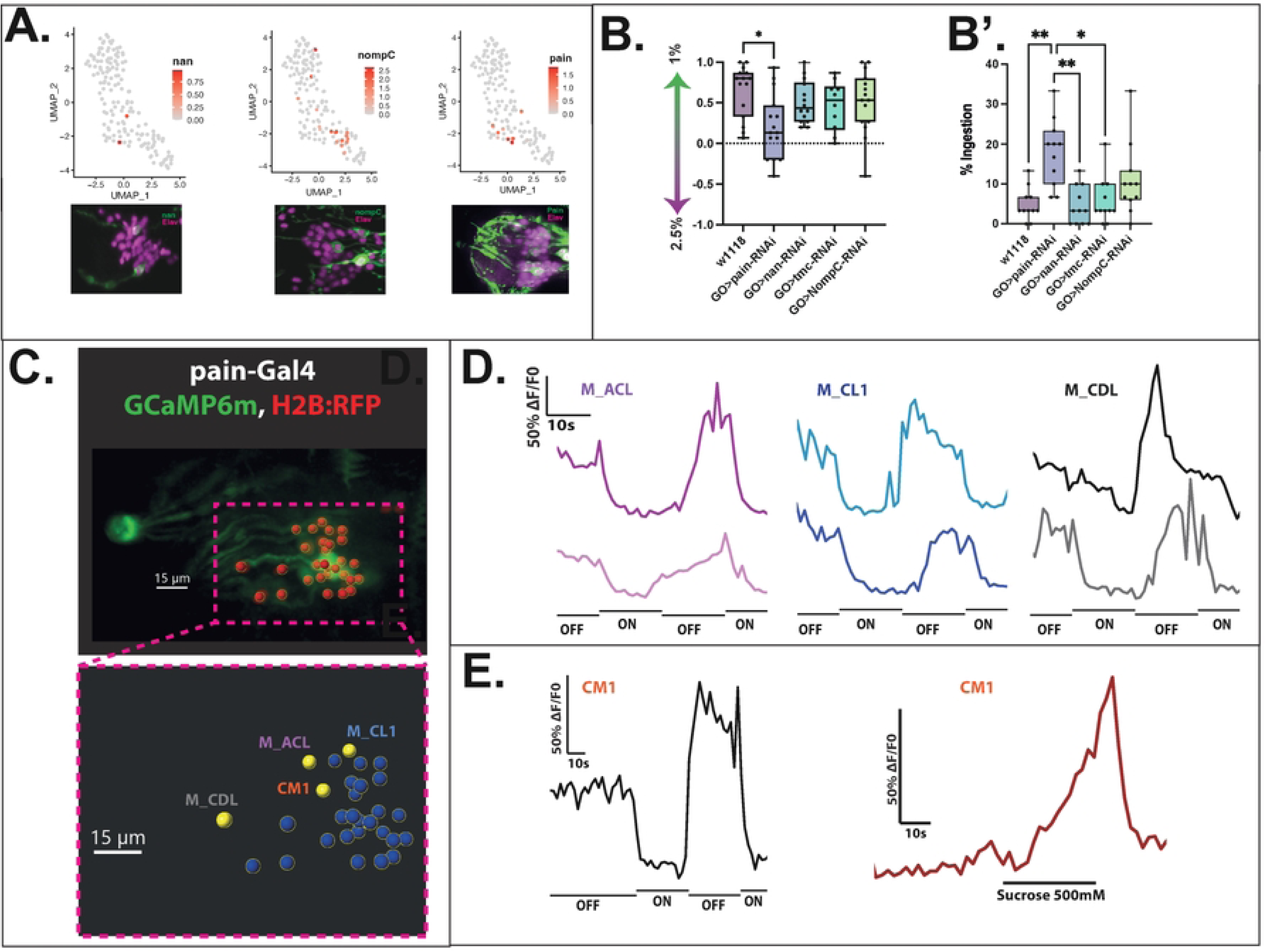
evaluation of genetic and physiological characteristics underlying mechanosensory function. A: Single-cell RNA sequencing and immunohistochemical stainings showing the expression of the mechanosensory genes *nanchug* (*nan*), *nompC*, and *painless* (*pain*). *nan* shows expression in 2 neurons of the head organs, *nompC* is expressed in 3 neurons, and a broad expression of *pain* can be observed. B: panel testing of putative mechanoreceptor genes shows that expressing *painless*, but not *nan*, *tmc*, or *nompC* RNAi results in a softness preference defect similar to complete Rpr-mediated ablation of GOs. B’: ingestion of blue-dyed hard agarose by *w*^1118^ larvae expressing total ablation of the GO (GO>rpr), as well as a set of mechanosensory RNAi-knockdown reporters. While control larvae (GO-GAL4 x *w*^1118^ and *w*^1118^ x UAS-rpr) do not ingest the hard agarose, larvae with ablated GO or larvae expressing *painless* RNAi in the GO show a greater degree of ingestion. N=10-15 for all behaviour assays. One-way ANOVA (Tukey’s multiple comparisons) test. **-p<0.01, ****-p<0.0001, ns-Not Significant (p>0.05). Not significant where not shown. C: Live calcium imaging view of 4 neurons (highlighted in yellow in panel C) which show relatively similar mechanosensory traces (D, E). One neuron, Centro-medial 1, shows responses to both mechanical stimulation and chemical stimulation (sucrose, 500mM) (E).

To more accurately pinpoint the principles of this texture sensing, we tested animals expressing RNAi for a range of known and putative mechanosensory genes, including those identified to be expressed in the peripheral chemosensory organs (Figure 3A). Here, we found that silencing expression of the TRPA family member *painless*, but none of the other candidates, results in a reduction of soft texture preference akin to silencing the entire organ (Figure 3B). Furthermore, we tested whether texture sensing contributes to ingestion decision-making by visually ascertaining the presence of blue-dyed agarose inside the animals. Previously, it was assumed that animals are unable, rather than unwilling, to ingest harder agarose as quickly as soft ^10^. However, we found that ablating the GOs increases immediate ingestion, and similarly to texture preference, driving *painless* RNAi results in a similar phenotype, whereby the animals more readily ingest harder agarose (Figure 3B’). This suggests that *painless* (*pain*)-expressing neurons play a part in informing the animal about the texture of the food.

### Identification of TOG neurons physiologically responding to mechanical stimulation

Using volumetric calcium imaging recordings, we tested whether applying a shear force by switching on and off of water flow through a microfluidic chamber (*i.e.*, applying pressure) elicits a response in *pain*-expressing neurons. Here, we found that 3-4 neurons in the TOG respond to this stimulus with a reduction of fluorescence, indicating a “silencing” effect of mechanical stimulation (Figure 3C, D). To test whether any of the responding neurons also carry a chemosensory role, we applied a sugar solution due to its broad and characterised response profile ^10^ to ascertain the presence or absence of multisensory responses. Intriguingly, we found that one neuron, which we named central-medial 1 (CM1) due to its anatomical position, responds to both mechanical stimulation and sucrose dynamically opposingly (Figure 3E).

Knowing the individual identity of the majority of the sucrose-sensitive neurons ^3,28^, we probed these neurons (C2, C5-7) using individual GAL-4 driver lines in an effort to identify CM1. While we did not observe mechanosensory responses in C2, C5, or C7, excitingly, we did observe a consistent response of C6, concordant with whole-organ imaging (Figure 4A, B).

**Figure 4:**
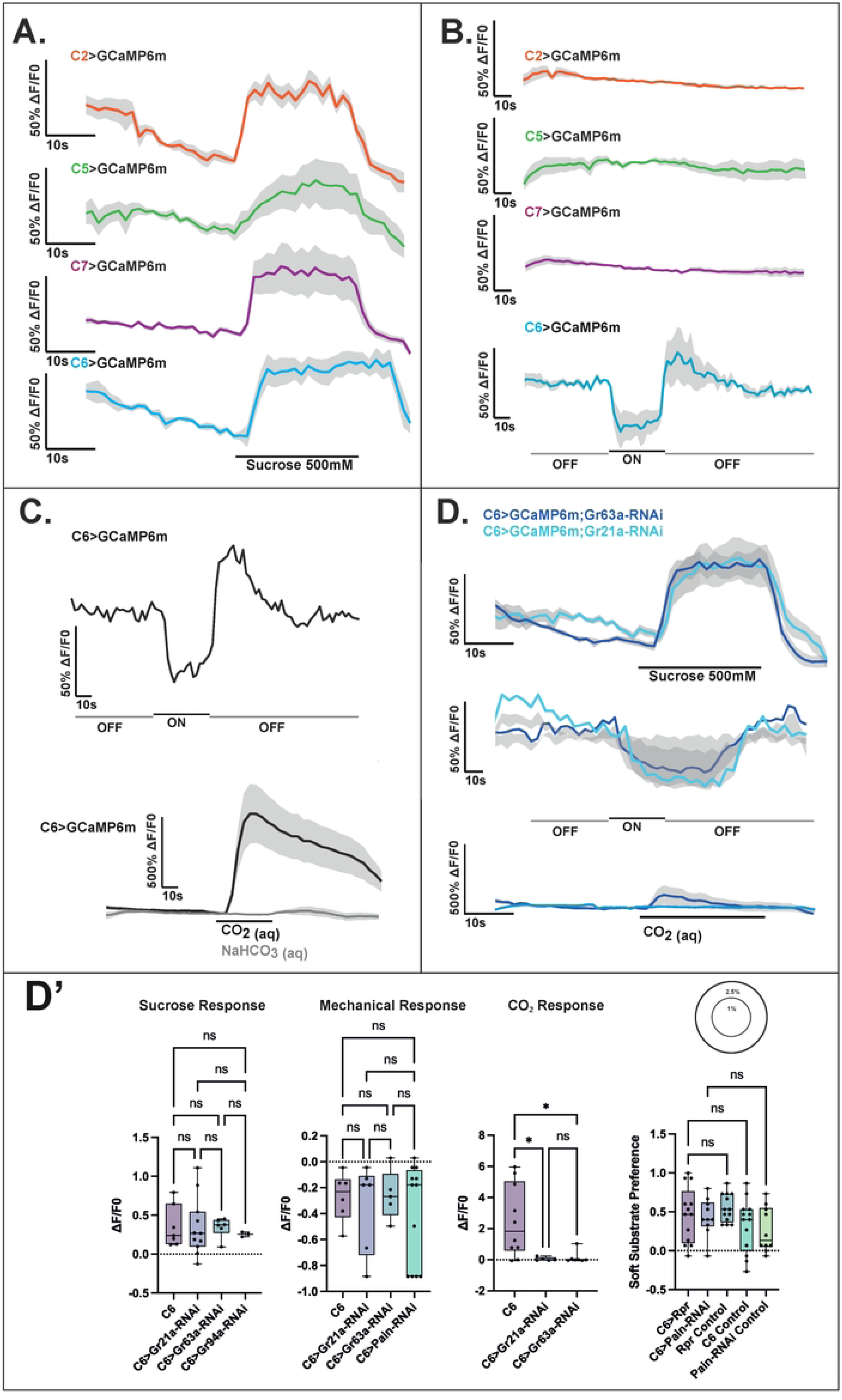
Identification of a multisensory and multimodal neuron in the GO. A: confirmation of single-neuron sucrose responses of comparable magnitude to CM1. C2, C5, C6, and C7 all show an approximately 50% fluorescence change when stimulated with sucrose. B: of the sucrose-sensitive neurons, only C6 shows a response to mechanical stimulation. C: responses of C6 to mechanical stimulation and carbonated water, in a live calcium imaging paradigm with representative traces shown. C6 displays a strong and consistent response to CO_2_, indicating that it bears a pseudo-olfactory role in carbon dioxide sensing. D: RNAi-mediated gene knockdown of carbon dioxide receptors *Gr21a* and *Gr63a* in C6 shows that both are required for carbonation sensing, but not required for sucrose or mechanosensory sensing (D’). N=10 for each bar, one-way ANOVA *-p<0.05, ns – not significant.

C6 is characterised by the individual GAL-4 drivers of the *Gr21a* and *Gr63a* receptors. Notably, these genes are known to be conserved across Diptera as essential to sensing carbon dioxide (CO_2_) ^17,29–31^. However, there has been no characterisation of physiological responses at the single neuron level in the larva. Using *Gr63a*-GAL4 as the driver for the expression of GCaMP6m, we performed recordings of the neurons before and during stimulation with aqueous CO_2_. Expectedly, we found a solid and robust activation response to CO_2(aq)_, but not to HCO ^-^ control, thus ensuring that the response is due to molecular CO_2_ and not to carbonate ions (Figure 3C).

Further, we confirmed that the mechanism for carbonation sensing relies on *Gr21a* and *Gr63a* through selective knockdown of expression using RNAi. Interestingly, in these conditions, the responses to sucrose and mechanical stimulation were not affected, suggesting independent mechanisms for the varied responses (Figure 4D, D’). Thus, we show that C6 is a multimodal chemosensory neuron that responds to both carbon dioxide and sucrose while it also exhibits responses to mechanical stimulation. In this light, C6 appears to be the first identified neuron which combines responses to multiple chemical and sensory modalities. In addition, as there is a noted lack of mechanical or sucrose phenotype of C6 under CO_2_-receptor knockdown, we propose that the mechanisms of physiological responses to tested stimuli are not linked to each other.

## Discussion

In this study, we determined a specific preference for food substrate texture in the *Drosophila* larva model. We also identify that the larva primarily employs the gustatory organs, akin to a human tongue, as key mechanosensors for texture evaluation. We also show that the commonly used agarose concentrations for larval behaviour correspond to more advanced fruit decomposition stages, which get softer with time. Although notable, this specific finding must be regarded as anecdotal due to the inability to control the genetics and harvest timing of fruits used. Nevertheless, this provides an insight into the physical properties of decomposing fruits, and how these properties relate to agarose substrates commonly used in behavioural experiments.

Next, we present evidence of multiple sensory modalities being coded in model gustatory organs (GOs). We show that GO ablation does not affect olfactory and light sensing but does affect taste and partial mechanical sensing. Additionally, we propose that the mechanoreceptor *painless*-expressing neurons affect the larva’s ability to make food choice decisions, both for navigation (seeking) and ingestion. Thus, we show that the mechanosensory neurons contained within GO are sufficient in sensing the mechanical properties of the food substrate, and the repression of mechanosensory genes in these cells is sufficient for creating food-choice decision defects. We also show that independent mechanisms (*i.e.* not *Gr21a*/*Gr63a*) contribute to physiological responses to mechanical stimulation.

While presumed before, the concept of mechanical perception being integrated within taste-sensing organs brings about fascinating questions about sensory integration in a numerically simple animal model. Finding that multimodal and multisensory neurons are present in the taste organs, we get further insight into the complexity of sensory processing ^32^. While the mechanisms and functions behind multisensory responses remain mostly elusive, we identify that they are at least partially independent, mimicking similar findings in *C. elegans* and adult *Drosophila* ^33–35^. Further screening of gene expression in individual multimodal and multisensory neurons across models is required to understand their full mechanisms. However, we believe that this is the first demonstration of mechanical and chemical perception within an individual sensory neuron via independent mechanisms in *Drosophila*.

Interestingly, promiscuity of mechanoreceptors such as the Transient receptor potential (TRP) family, where a wide variety of functions have been observed, from nociception to thermosensation across different models, may play a role in the varied mechanosensitive responses observed here ^36–38^. Additionally, the co-expression of different mechanosensory genes, including the TRP family, within sensory neurons is also described ^28,32,34^, which, in coordination with our results, reveals an intriguing path for investigating individual receptor roles in the physiological responses of GSNs.

Moreover, these findings allow us to ask in-depth questions about the central processing of taste stimuli. It is possible to speculate that rather than transmitting information about an “appetitive” or “aversive” stimulus by individual neurons, as is the case for olfaction, the brain integrates the signals from the whole taste system before making decisions. That is, as multiple neurons sense the same stimuli, and yet the multimodal combinations are different, this creates a large sensory range when considering the number of unique combinations of neuronal responses. Further, the recently-released connectome dataset can allow for studies of local processing within the primary gustatory neuropil – sub-esophageal zone (SEZ), which, coupled with recent discoveries about peripheral sensory physiology as in this study, can shed more light on the logic of taste processing in the *Drosophila* larva and beyond. For instance, one question that can be asked is whether different input modalities result in different signal outputs. For example, connectomic studies suggest that sensory neurons communicate with one another via axon-axonal interactions before they reach the brain or may result in outputs to synapses at different brain targets ^39^. This could allow, for example, a neuron to modulate the signals perceived by the brain from its neighbours, which may, in turn, explain the reason for multimodality in individual gustatory sensory cells.

## Materials and Methods

### Fly stocks and husbandry

Flies used for experimental crosses were maintained at 25°C on a 12/12 hour dark-light cycle. Fly stocks were fed standard cornmeal food. The following lines were used in the study, including from the Bloomington Drosophila Stock Centre (BDSC) and the Vienna Drosophila Resource Centre (VDRC):

**Supplementary table 1:**
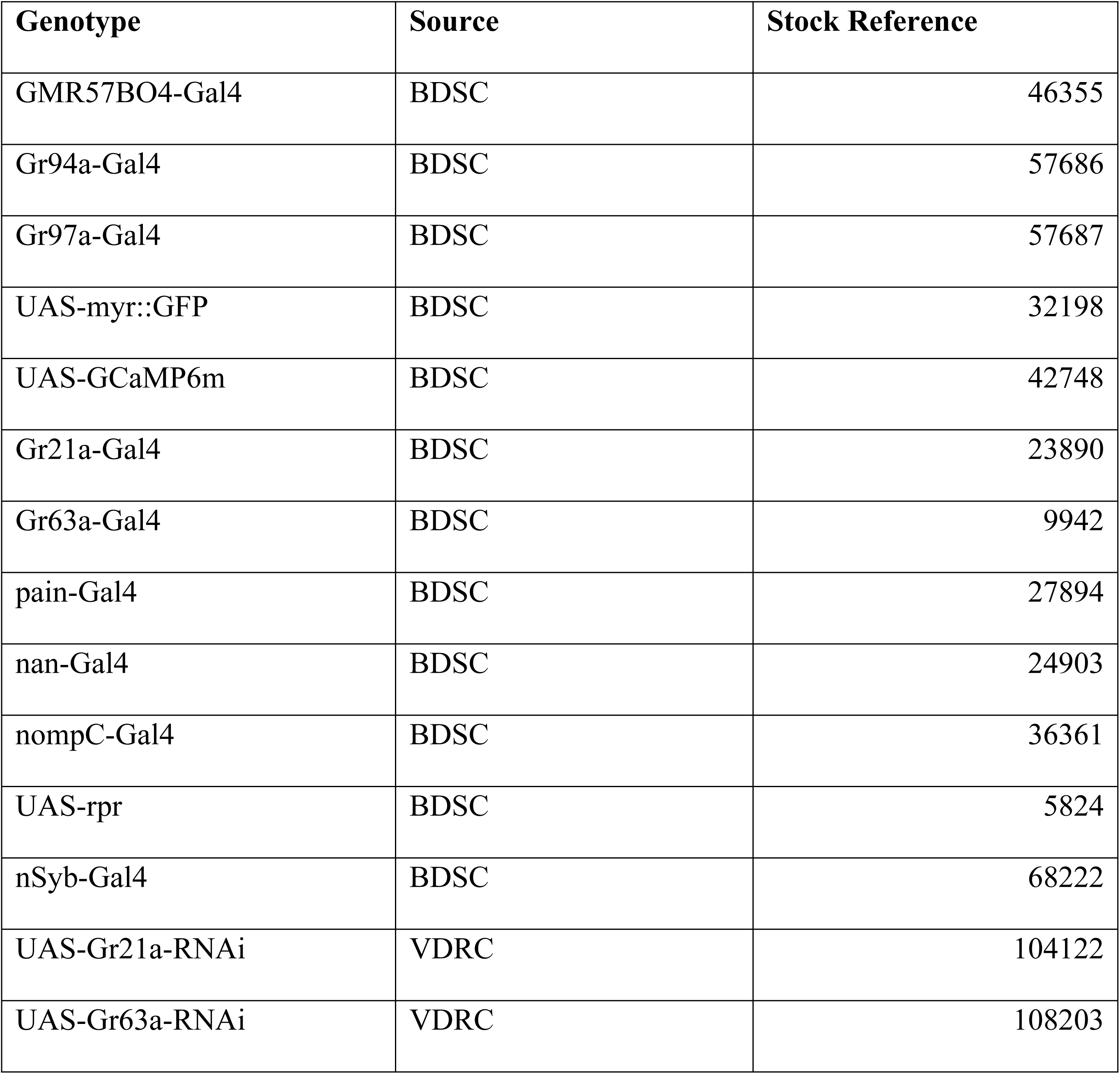
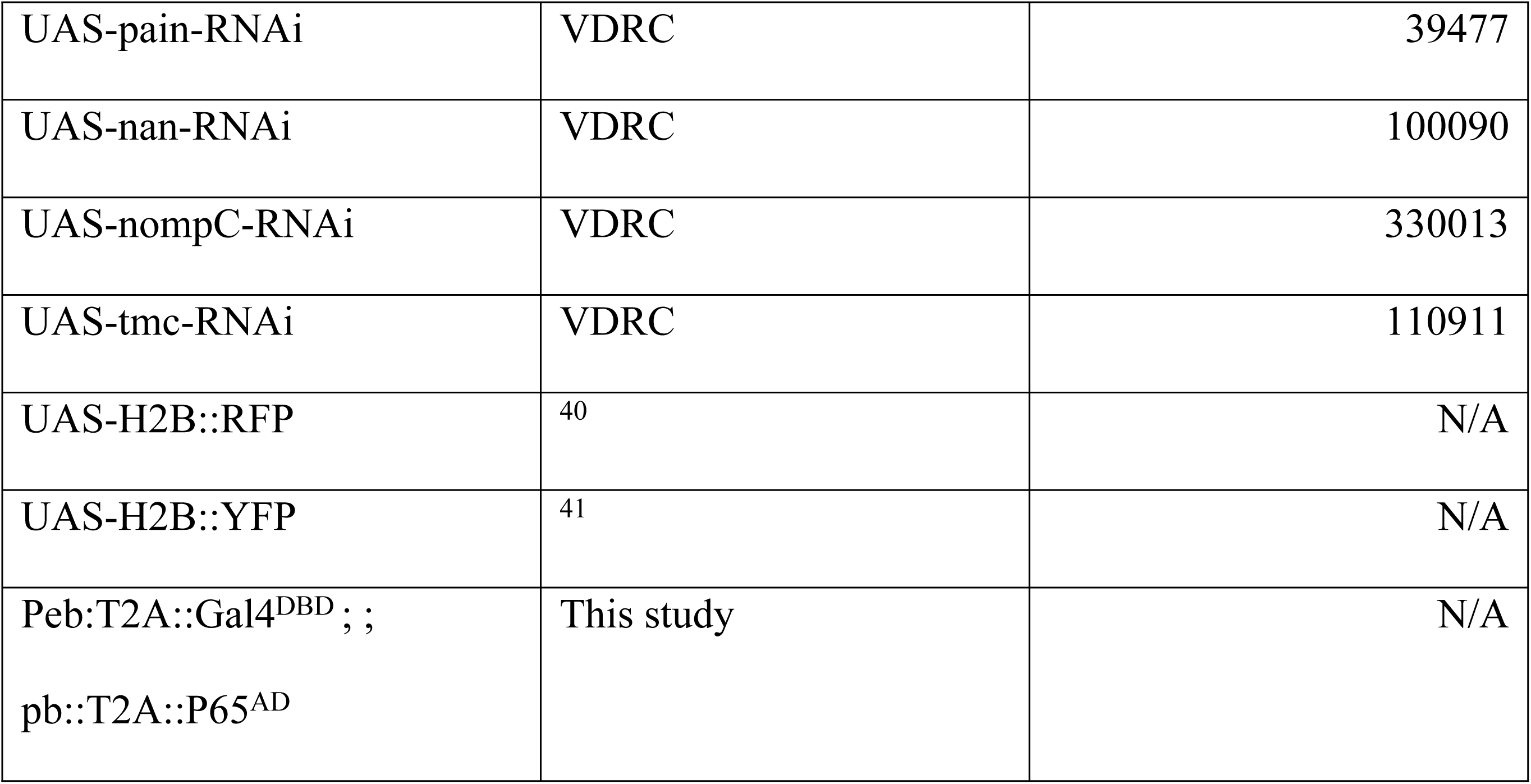
Fly stocks used.

### Generation of a split-GAL4 line for gustatory organs

In order to generate a GO-specific split-GAL4 line, we chose two genes that are co-expressed in the cells of the taste organs but do not show overlapping expression in other tissues. We opted for the multiple zinc-finger transcription factor Pebbled (Peb) and the homeodomain transcription factor Proboscipedia (Pb) (Figure 1C). We decided to produce a split-GAL4 line expressing the DNA-binding domain (DBD) of GAL4 under the control of *peb* regulatory sequences and the p65 (GAL4) activation domain (AD) under the control of *pb* regulatory sequences. To ensure that the split-GAL4 constructs are expressed in the same cells as the endogenous genes, we fuse them in frame to the coding sequences of *peb* and *pb* using the CRISPR-Cas9 technique. To maintain the function of the endogenous transcription factors Peb and Pb and the split-GAL4 protein parts, the two proteins were connected with an autocatalytic peptide (T2A). After translation of the fusion proteins, the 18 amino acid-long T2A sequence will cleave itself just before its last amino acid, separating the endogenous protein from the attached split-GAL4 fragment and allowing both proteins to function independently. A protein zipper domain will combine both domains to a functional GAL4 complex in cells that express both fusion proteins. The *GAL4 DBD* fragment and zipper domain were PCR amplified from plasmid *pBPZpGAL4DBDUw* (Pfeiffer et al., 2010, Rubin Lab, addgene No. 26233) and cloned into *pBluescript*. The *T2A* sequence from plasmid *pF3BGX-T2A-p65-AD* ^43^, Shu Kondo lab, addgene No. 138395) was added to the *zipper-GAL4DBD*. An 837 bp fragment of the C-terminal end of *peb* and 1066 bp of its 3’UTR were PCR amplified from genomic DNA isolated from *nos-Cas9* flies to serve as homology arms for the CRISPR template. Since the first CRISPR site used for insertion of the *GAL4* fragment is located within the *peb* coding sequence, the last 12 amino acids of Peb will be replaced with the T2A peptide after autocatalytic cleavage of the Peb-GAL4DBD fusion protein. The *p65AD* fragment, including *T2A* peptide, *NLS* and protein *zipper* domain, was PCR amplified from plasmid *pF3BGX-T2A-p65-AD* and cloned into *pBluescript*. A1434 bp fragment containing the last intron and last exon of pb and a 1155 bp fragment containing its 3’UTR and downstream genomic sequence, PCR amplified from *nos-Cas9* genomic DNA, were added as homology arms for the CRISPR template. In this construct the entire Pb protein was fused to the T2A-NLS-GAL4 fragment, so that after autocatalytic cleavage the T2A peptide will be attached to the last amino acid of the Pb protein.

The *peb-GAL4DPD* template was injected into embryos of flies expressing Cas9 under the control of the *nanos* promoter (Bloomington stock no. 54591) along with a *pCFD4-U6:1_U6:3tandemgRNAs* plasmid (Port et al., 2014, Simon Bullock Lab, addgene no. 49411) expressing two gRNAs for CRISPR sites located at the end of the *peb* coding sequence and in its 3’UTR. The *pb-p65AD* template was co-injected with a *pCFD4-U6:1_U6:3tandemgRNAs* plasmid expressing two gRNAs for CRISPR sites located in the 3’UTR of *pb*. After eclosion, the *peb-GAL4DBD*-injected flies were crossed to a first chromosome balancer line (*N/FM7C-GFP*) and the *pb-p65AD*-injected flies were crossed to a third chromosome balancer line (*w;; Dr, e/TM3*). Single F1 offspring flies were crossed again with the appropriate balancer lines and PCR-screened for the presence of the *GAL4*-fragments. For each GAL4 fragment, two independent insertion lines were established. The two split-*GAL4* fragments inserted on chromosomes 1 and 3 were combined in a single line.

**Supplementary table 2:**
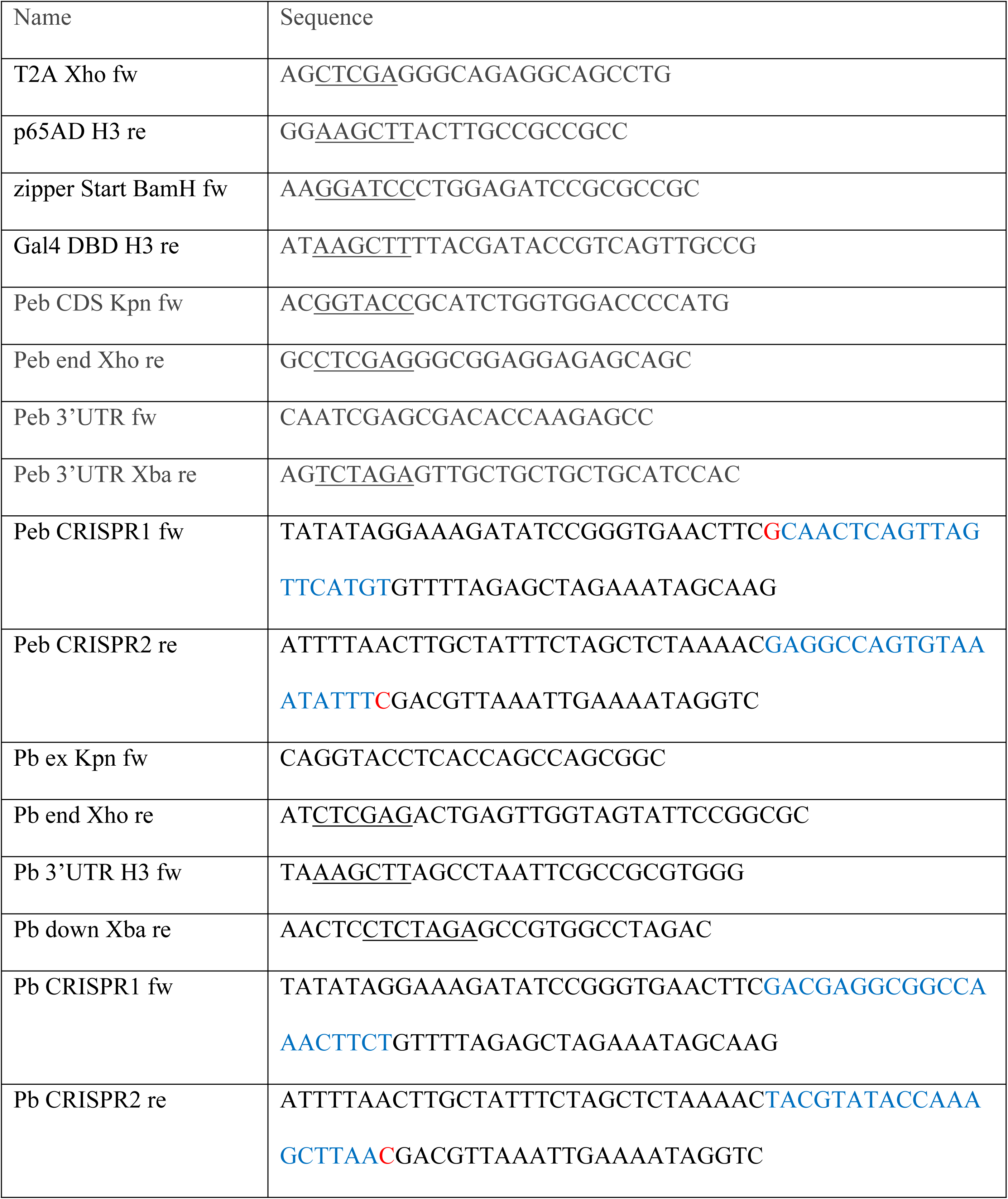
Primers used for generation of the split-*GAL4* lines (restriction sites underlined, CRISPR sites blue)

### Larval behaviour

3^rd^ instar, pre-wandering stage larvae were collected 72-96 hours following crossing. Crosses were made on standard cornmeal food supplemented with liquid yeast paste. Animals were kept at 25°C on a 12h/12h dark-light cycle.

#### Larval behaviour – two choice assays

Two-choice assays were performed as described thoroughly in Maier et al., 2021, with the following modifications:

Hardness preference: 94mm petri dishes were filled with either 0.1%, 1%, or 2.5% agarose boiled in nanopore water until dissolved. For 0.1% vs 1% assays, a 66mm central circular cut-out was filled with the other concentration, respectively. A similar preparation was made for the 1% vs 2.5% assays. Plates containing the harder substance in the middle vs outside were randomly selected for each test. Thirty larvae were collected and rinsed in tap water before being placed on the centre of each plate. The number of larvae on each substrate was recorded at 2, 5, and 15 minutes. For larvae crossing the edge of the two substrate concentrations, the affirmative decision about the condition was made depending on the location of the mouth hooks. Following this, a preference index (PI) was calculated using the following formula:

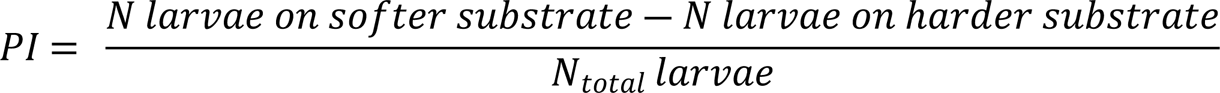

Light preference: 94mm petri dishes were filled with 2.5% agarose. Thirty 3^rd^ instar larvae per trial (crossed, maintained, and collected as above) were placed in the middle of the dish. Half of the dish lid was covered with aluminium foil, and the preparation was illuminated from a projector (Epson LCD Projector model H763B, default settings) positioned 70cm above the experimental space, emitting a white (RGB: 255, 255, 255) light. The number of larvae on the illuminated side was counted, and a preference index was calculated as follows to avoid exposure of the dark side to light:

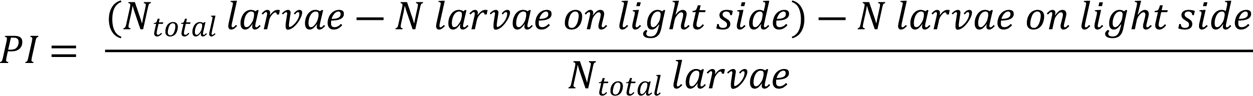

#### Larval behaviour – ingestion assay

Petri dishes were filled with 2.5% agarose supplemented with Brilliant Blue dye (2% w/v). Thirty 3^rd^ instar larvae per trial (crossed, maintained, and collected as above) were placed on the blue agarose and left to wander for 2 minutes. Larvae were then collected, briefly washed in tap water, and examined for the presence of blue dye in the digestive tract, indicating ingestion.

#### Live calcium imaging

Live calcium imaging was performed as described in van Giesen et al., 2016b and analysed as described in Maier et al., 2021. In short, L3 larval heads were dissected posteriorly to the brain and mounted inside a microfluidics chamber, and sealed in with 2% agarose in AHL saline. Water was pumped through the chip for the first 60s of the recording, followed by a 30s stimulation with tastant and a 30s water wash. The following adjustments were made to the protocol:

Mechanical stimulation: to simulate mechanical pressure, larval heads were positioned within the microfluidic device and briefly washed with millipore water, with the flow being switched off 5s before the start of the recording, with the larval head remaining in the aqueous environment of the microfluidic chamber. Macros were adjusted to switch the water flow on at the 60s time point. Thus, corresponding neuronal responses were interpreted to result from shear stress (mechanical stimulation) rather than hygrosensation ^46,47^.

#### Immunohistochemistry

Embryos: embryo collection, dechorionation, and fixation were performed as described in Müller, 2008, with the 3.7% formaldehyde (Merck, 1.04003.1000) fixation method being employed before antibody incubation. Primary antibodies used: Rat-anti-Elav (DSHB 7E8A10, 1:50 dilution), Chicken-anti-GFP (Abcam 13970, 1:2000 dilution), Mouse-anti-Pebbled (DSHB 1G9, 1:50 dilution), Rabbit-anti-DsRed (TaKaRa 632496, 1:1000 dilution), Rabbit-anti-Pb (Cribbs, 1992 ^49^, 1:100 dilution). Fixed embryos were rehydrated in 1X-PBS (neoFroxx, 1346LT050) and briefly washed before incubation with primary antibodies in 1X-PBS with 0.3% Triton-X-100 (Roth, 3051.1) (PBS-T), overnight at 4°C. The following day, the primary antibodies were removed, and the embryos were washed at RT with PBS-T for at least two hours, replacing the PBS-T every 30 minutes. Secondary antibodies used: Donkey-anti-Mouse Alexa 488 (Molecular Probes A21202, 1:1000 dilution), Donkey-anti-Rat Alexa 647 (Jackson ImmunoResearch 712-605-150 1:1000 dilution), Donkey-anti-Rabbit Alexa 647 (Molecular Probes A31573, 1:1000 dilution), Goat-anti-Chicken Alexa 488 (Molecular Probes A11039, 1:1000 dilution). Washed embryos were then incubated with the appropriate secondary antibodies in PBS-T overnight at 4°C. The following day, the secondary antibody solution was removed, and embryos were briefly washed with PBS-T 3 times, before being washed in PBS-T with DAPI (Roth, 6335.2) (1:50000 dilution) for 30 minutes at RT. The DAPI was then removed, and the samples were washed in PBS-T for at least 2 hours, changing the PBS-T every 30 minutes. Following this, the PBS-T was replaced with mounting medium (90% Glycerol (Fischer Scientific BP229-1), 0.5% N-propyl gallate (Sigma P3130), 20 mM Tris (Fischer Scientific BP152-5, pH 8.0) for at least 1 hour at RT before mounting on standard glass slides.

L3 Larvae: Larval heads were dissected in PBS anterior to the mouth hooks, removing as much cuticle as possible without compromising the structural integrity of the samples. Dissected samples were kept in PBS on ice until the fixation step (dissection time should not exceed 1 hour). Samples were then fixed in 3.7% formaldehyde in PBS for a minimum of 18 and a maximum of 30 minutes, shaking at RT. The formaldehyde solution was then removed, and the heads were briefly rinsed with PBS-T before being washed in PBS-T for at least 2 hours, replacing the PBS-T every 30 minutes. Following this, the immunohistochemistry steps do not differ from those described for embryo labelling above. For improved penetration of the antibodies, 0.5% Triton X-100 was used.

Imaging and processing: microscopy was carried out on the Leica Stellaris 8 Falcon confocal microscope, using the Plan APO 40x/1.10 water immersion objective. Acquired images were processed using Fiji ImageJ, and Figures arranged with Adobe Illustrator.

#### Single-cell chemosensory cell suspension preparation for DisCo

3^rd^ instar larvae of genotype *nsyb*-Gal4 > UAS-*mcd8::GFP; Or83b::RFP* were collected from food, washed in tap water, PBS, dropped in ethanol and again PBS and dissected in ice-cold PBS in such a manner that only the external chemosensory organs were kept, avoiding to include also the pharyngeal tissue containing internal chemosensory organs. The isolated material was placed on ice in elastase 1 mg/ml in siliconised 2-ml tubes. After dissecting 20-30 larvae (20-25 minutes), the tube sample was placed at room temperature to initiate digestion. After 30 minutes, the tissue was washed in PBS+BSA0,05% and dissociated by up-down pipetting 120 times using ssiliconised 200p pipette tips. Separated TOG and DOG (expressing *Or83b*::*RFP*) organs were detected using a fluorescence stereomicroscope and manually picked with a glass micropipette, placed in a final dissociation enzyme mix of Collagenase 1mg/ml + Elastase 0.5mg/ml for 10-15 minutes for single-cell suspension. The reaction was stopped with PBS + BSA 0,05%. Murine inhibitor was added at each step of the dissociation protocol.

#### Deterministic co-encapsulation (DisCo) of chemosensory neurons for single cell transcriptomics

Microfluidic chip design, fabrication and device handling are described elsewhere ^24^. Following organ dissociation, target cell suspension was diluted in the cell loading buffer containing PBS 0.01 % BSA (Sigma B8667), 6% Optiprep (Sigma D1556) and Murine RNase inhibitor (NEB M0314L) in the loading tip connected to the DisCo chip. After bead-cell in droplet co-encapsulation, sample droplets were transferred to a bead collection chip. Subsequently to bead capture, washing, reverse transcription (Thermo Scientific EP0753) and Exonuclease I (NEB M0293L) reactions were performed on-chip ^50^. Beads containing cDNA were then eluted, and cDNA was amplified for 21 cycles using Kapa HiFi Hot start ready mix (Roche #07958935001). Amplified cDNA was then purified (GC biotech CPCR-0050) for quality assessment with Fragment Analyzer (Agilent). Libraries were then tagmented using in-house Tn5 ^51^, size selected and purified for sequencing on NextSeq 500 system (Illumina) following recommendations from original Drop-seq protocol ^52^ (20 bp for read 1 and 50 bp for read2) at sequencing depth above 400.000 reads per cell.

#### Single-cell data pre-processing and analysis

The data analysis was performed using the Drop-seq tools package ^52^. After pre-processing, reads were aligned to *Drosophila melanogaster* reference genome (Ensembl version 86) using STAR (version 2.7.0.e) ^53^. Following the alignment, BAM files were processed using the initial package and read-count matrices were generated.

Downstream analysis was done using the Seurat package ^26^ version 3.1.2 in R version 4.2.2, in Rstudio version 2022.12.0+353. Individual data sets were loaded and used to create separate snormalised and scaled Seurat objects of minimum 400 genes per cell. In order to apply unique cell filters, we merged the data and then excluded cells with high gene numbers and high UMIs as potential doublets and cells with high mitochondrial gene percentages indicating potential apoptotic cells. Due to the observed correlation between cells with high gene number (nGene) and cells with high UMIs (nCount), by applying UMI threshold at 50000 we also eliminated cells with more than 4000 genes. Cells with mitochondrial gene percentages under 9% were kept for further analysis. Data was then integrated to circumvent batch effects using Seurat functions *FindIntegrationAnchors* and *IntegrateData*. As we were interested in scharacterising neurons, we used the *subset* function to keep only cells expressing *nSyb* or *peb* neuronal markers, excluding eventual surrounding tissue or cuticle cells. On the final dataset of 153 neurons PCA (principal component analysis) computation was followed by UMAP embedding, and clustering was performed at 0.5 resolution.

#### Statistical analysis

Statistical testing and visualisation of data pertaining to live calcium imaging and single-cell sequencing was performed with R version 4.2.2 in R Studio version 2022.12.0+353. Quantitative analysis of behaviour and ΔF/F0 values was performed using Prism 9 (GraphPad Software), with bars showing min-max, with all points shown. The type of test, p values, and sample size for each graph are provided in the respective Figure legends. Significance is displayed as follows: ns – not significant, *P<*0.05(*), *P*<0.01(**), *P*<0.001(***), *P*<0.0001(****). Figures were assembled using Adobe Illustrator 2024, including the use of the integrated generative AI feature for the creation of fruit icons in Figure 1.

#### Substrate hardness evaluation

Compression tests were performed using an Anton Paar MCR 702 rheometer equipped with a CTD600 convection temperature device. A plate-plate geometry with an 8mm diameter and a compression speed of 100 um/s was used. All measurements were carried out at room temperature. 8mm disks were cut from the samples and placed between the plates, applying a normal force of 0.1 N to ensure good sample loading.

Agarose plates were prepared in 50ml of water at concentrations of 0.1% (gelling point), and in the range from 0.25% to 2.5% (w/v) in 0.25% increments by boiling the solutions for 2 minutes, with the volume adjusted to 50ml post-boiling.

For fruits, five individual units of pear, apple, pineapple, and banana were cut into sections of ∼5mm thickness and then stored in a humidified incubator (100% RH, 25°C) to simulate the decomposition process. For five consecutive days, 8mm disks were cut from the sections and immediately measured. The compression modulus was then calculated from the initial slope of the obtained stress-strain curves. Three measurements were repeated for each sample.

## Acknowledgments

We would like to express our gratitude to the Swiss National Science Foundation (SNSF) for providing the funding for this project (grant 310030_219348 and IZKSZ3_218514 to SGS). We are also thankful to the University of Fribourg (UniFr) and the members of the Department of Biology for their support and help throughout our research. We especially thank Boris Egger and Felix Meyenhofer of the UniFr Bioimage Core Facility for their immense support in the imaging. We thank the Bloomington Drosophila Stock Centre and the Vienna Drosophila Resource Centre, Boris Egger and Shahnaz Lone for sourcing and sharing fly stocks.

We also thank all members of the Sprecher Lab for their invaluable feedback on our experimental design, procedures, and reporting. Their insights and suggestions are invaluable to our work.

Anton Paar GmbH is gratefully acknowledged for the loan of the MCR702 MultiDrive rheometer and their excellent technical support.

## Declaration of generative AI and AI-assisted technologies

During the preparation of this work, the authors employed the AI image generation feature in Adobe illustrator to create minor fruit graphics present in Figure 1. The authors have reviewed and edited the content as needed and take full responsibility for the published material.

## Author contributions

NK, LM, SGS designed, performed, and analysed calcium imaging experiments.

LM, JB, MB, CBA, BD, SGS performed scRNAseq experiments and analysis.

LM, NK, JYK, BD, SGS designed and coordinated experiments.

CBA, LM identified GO markers.

CF created transgenic lines used in this study.

NK, SGS designed, performed, and analysed behavioural experiments.

NK, CBA, LM conducted immunofluorescence expression analysis.

All authors contributed to writing and editing of the manuscript.

---

**Figure S1: single-cell sequencing procedure**. A: position of TOG and DOG within the larval head. B: isolation of TOG cells by fluorescent discrimination of GFP expressed in the pattern of the *Ir76b* promoter (subset of TOG neurons), and DOG through expression of Or83b::RFP (olfactory sensory neurons). B’: filtering of the cells by neuronal markers *Neuroglian* (*Nrg*), *Synaptobrevin* (*Syb*), *neuronal Synaptobrevin* (*nSyb*), and *pebbled* (*peb*). C: identification of olfactory neurons (marked by *Orco* expression) and taste neurons (marked by *Pb*) expression, showing no visible overlap of the two genes, as confirmed by immunohistochemical analysis.

**Figure S2: expression analysis of different receptor types in the larval head organs**. A: identified *Gr* genes. B: co-expression of specific *Gr* genes allows for identification of individual neurons, indicated by arrows. C: putative mechanoreceptor gene *pain* is co-expressed with *Gr66a* in at least one cell (arrow). D: identified *Or* genes. E: identified *Ir* genes. F: identified *Ppk* family genes.

## References

1. Oh, S. M., Jeong, K., Seo, J. T. & Moon, S. J. Multisensory interactions regulate feeding behavior in Drosophila. Proceedings of the National Academy of Sciences 118, e2004523118 (2021).

2. Apostolopoulou, A. A., Rist, A. & Thum, A. S. Taste processing in Drosophila larvae. Front. Integr. Neurosci. 9, (2015).

3. Rist, A. & Thum, A. S. A map of sensilla and neurons in the taste system of drosophila larvae. Journal of Comparative Neurology 525, 3865–3889 (2017).

4. Kim, D., Alvarez, M., Lechuga, L. M. & Louis, M. Species-specific modulation of food-search behavior by respiration and chemosensation in Drosophila larvae. eLife 6, e27057 (2017).

5. Kudow, N., Kamikouchi, A. & Tanimura, T. Softness sensing and learning in Drosophila larvae. Journal of Experimental Biology 222, jeb196329 (2019).

6. Liman, E. R., Zhang, Y. V. & Montell, C. Peripheral Coding of Taste. Neuron 81, 984–1000 (2014).

7. Thorne, N., Chromey, C., Bray, S. & Amrein, H. Taste Perception and Coding in Drosophila. Current Biology 14, 1065–1079 (2004).

8. Yarmolinsky, D. A., Zuker, C. S. & Ryba, N. J. P. Common Sense about Taste: From Mammals to Insects. Cell 139, 234–244 (2009).

9. Caicedo, A., Kim, K.-N. & Roper, S. D. Individual mouse taste cells respond to multiple chemical stimuli. J Physiol 544, 501–509 (2002).

10. Maier, G. L., Komarov, N., Meyenhofer, F., Kwon, J. Y. & Sprecher, S. G. Taste sensing and sugar detection mechanisms in Drosophila larval primary taste center. eLife 10, e67844 (2021).

11. van Giesen, L. et al. Multimodal stimulus coding by a gustatory sensory neuron in Drosophila larvae. Nature Communications 7, 10687 (2016).

12. Stocker, R. F. Design of the Larval Chemosensory System. in Brain Development in Drosophila melanogaster (ed. Technau, G. M.) 69–81 (Springer, New York, NY, 2008). doi:10.1007/978-0-387-78261-4_5.

13. Komarov, N. & Sprecher, S. G. The chemosensory system of the Drosophila larva: an overview of current understanding. Fly 16, 1–12 (2022).

14. Hanna, L., et al. Evaluating old truths: Final adult size in holometabolous insects is set by the end of larval development. Journal of Experimental Zoology Part B: Molecular and Developmental Evolution 340, 270–276 (2023).

15. Vosshall, L. B. & Stocker, R. F. Molecular Architecture of Smell and Taste in *Drosophila*. Annu. Rev. Neurosci. 30, 505–533 (2007).

16. Mishra, D. et al. The molecular basis of sugar sensing in Drosophila larvae. Curr Biol 23, 1466–1471 (2013).

17. Jones, W. D., Cayirlioglu, P., Grunwald Kadow, I. & Vosshall, L. B. Two chemosensory receptors together mediate carbon dioxide detection in Drosophila. Nature 445, 86–90 (2007).

18. Alves, G., Sallé, J., Chaudy, S., Dupas, S. & Manière, G. High-NaCl Perception in Drosophila melanogaster. J. Neurosci. 34, 10884–10891 (2014).

19. Apostolopoulou, A. A., Mazija, L., Wüst, A. & Thum, A. S. The neuronal and molecular basis of quinine-dependent bitter taste signaling in Drosophila larvae. Front. Behav. Neurosci. 8, (2014).

20. Delventhal, R. & Carlson, J. R. Bitter taste receptors confer diverse functions to neurons. eLife 5, e11181 (2016).

21. Liu, L. et al. Contribution of Drosophila DEG/ENaC Genes to Salt Taste. Neuron 39, 133– 146 (2003).

22. DeGennaro, M. et al. orco mutant mosquitoes lose strong preference for humans and are not repelled by volatile DEET. Nature 498, 487–491 (2013).

23. Larsson, M. C. et al. Or83b Encodes a Broadly Expressed Odorant Receptor Essential for Drosophila Olfaction. Neuron 43, 703–714 (2004).

24. Bues, J. et al. Deterministic scRNA-seq captures variation in intestinal crypt and organoid composition. Nat Methods 19, 323–330 (2022).

25. Satija, R., Farrell, J. A., Gennert, D., Schier, A. F. & Regev, A. Spatial reconstruction of single-cell gene expression data. Nat Biotechnol 33, 495–502 (2015).

26. Stuart, T. et al. Comprehensive Integration of Single-Cell Data. Cell 177, 1888–1902.e21 (2019).

27. Rusch, D. B. & Kaufman, T. C. Regulation of proboscipedia in Drosophila by Homeotic Selector Genes. Genetics 156, 183–194 (2000).

28. Kwon, J. Y., Dahanukar, A., Weiss, L. A. & Carlson, J. R. Molecular and Cellular Organization of the Taste System in the Drosophila Larva. J. Neurosci. 31, 15300–15309 (2011).

29. Kumar, A. et al. Contributions of the Conserved Insect Carbon Dioxide Receptor Subunits to Odor Detection. Cell Reports 31, 107510 (2020).

30. Kwon, J. Y., Dahanukar, A., Weiss, L. A. & Carlson, J. R. The molecular basis of CO2 reception in Drosophila. PNAS 104, 3574–3578 (2007).

31. Robertson, H. M. & Kent, L. B. Evolution of the Gene Lineage Encoding the Carbon Dioxide Receptor in Insects. Journal of Insect Science 9, 1–14 (2009).

32. Sánchez-Alcañiz, J. A. & Benton, R. Multisensory neural integration of chemical and mechanical signals. BioEssays 39, 1700060 (2017).

33. Biron, D., Wasserman, S., Thomas, J. H., Samuel, A. D. T. & Sengupta, P. An olfactory neuron responds stochastically to temperature and modulates Caenorhabditis elegans thermotactic behavior. Proceedings of the National Academy of Sciences 105, 11002– 11007 (2008).

34. Jeong, Y. T. et al. Mechanosensory neurons control sweet sensing in Drosophila. Nat Commun 7, 12872 (2016).

35. Mellem, J. E., Brockie, P. J., Zheng, Y., Madsen, D. M. & Maricq, A. V. Decoding of Polymodal Sensory Stimuli by Postsynaptic Glutamate Receptors in C. elegans. Neuron 36, 933–944 (2002).

36. Bellemer, A. Thermotaxis, circadian rhythms, and TRP channels in Drosophila. Temperature 2, 227–243 (2015).

37. Julius, D. TRP Channels and Pain. Annual Review of Cell and Developmental Biology 29, 355–384 (2013).

38. Kashio, M. & Tominaga, M. TRP channels in thermosensation. Current Opinion in Neurobiology 75, 102591 (2022).

39. Miroschnikow, A. et al. Convergence of monosynaptic and polysynaptic sensory paths onto common motor outputs in a Drosophila feeding connectome. eLife 7, (2018).

40. Langevin, J. et al. Lethal Giant Larvae Controls the Localization of Notch-Signaling Regulators Numb, Neuralized, and Sanpodo in Drosophila Sensory-Organ Precursor Cells. Current Biology 15, 955–962 (2005).

41. Bellaïche, Y., Gho, M., Kaltschmidt, J. A., Brand, A. H. & Schweisguth, F. Frizzled regulates localization of cell-fate determinants and mitotic spindle rotation during asymmetric cell division. Nat Cell Biol 3, 50–57 (2001).

42. Pfeiffer, B. D. et al. Refinement of Tools for Targeted Gene Expression in Drosophila. Genetics 186, 735–755 (2010).

43. Kondo, S. et al. Neurochemical Organization of the Drosophila Brain Visualized by Endogenously Tagged Neurotransmitter Receptors. Cell Reports 30, 284–297.e5 (2020).

44. Port, F., Chen, H.-M., Lee, T. & Bullock, S. L. Optimized CRISPR/Cas tools for efficient germline and somatic genome engineering in Drosophila. Proceedings of the National Academy of Sciences 111, E2967–E2976 (2014).

45. van Giesen, L., Neagu-Maier, G. L., Kwon, J. Y. & Sprecher, S. G. A microfluidics-based method for measuring neuronal activity in Drosophila chemosensory neurons. Nature Protocols 11, 2389–2400 (2016).

46. Brown, T. D. Techniques for mechanical stimulation of cells in vitro: a review. Journal of Biomechanics 33, 3–14 (2000).

47. Gong, J. et al. Shear stress activates nociceptors to drive Drosophila mechanical nociception. Neuron (2022) doi:10.1016/j.neuron.2022.08.015.

48. Müller, H.-A. J. Immunolabelling of embryos. Methods Mol Biol 420, 207–218 (2008).

49. Cribbs, D. L., Pultz, M. A., Johnson, D., Mazzulla, M. & Kaufman, T. C. Structural complexity and evolutionary conservation of the Drosophila homeotic gene proboscipedia. EMBO J 11, 1437–1449 (1992).

50. Biočanin, M., Bues, J., Dainese, R., Amstad, E. & Deplancke, B. Simplified Drop-seq workflow with minimized bead loss using a bead capture and processing microfluidic chip. Lab on a Chip 19, 1610–1620 (2019).

51. Picelli, S. et al. Tn5 transposase and tagmentation procedures for massively scaled sequencing projects. Genome Res. 24, 2033–2040 (2014).

52. Macosko, E. Z. et al. Highly Parallel Genome-wide Expression Profiling of Individual Cells Using Nanoliter Droplets. Cell 161, 1202–1214 (2015).

53. Dobin, A. et al. STAR: ultrafast universal RNA-seq aligner. Bioinformatics 29, 15–21 (2013).

